# Quantifying the tradeoff between sequencing depth and cell number in single-cell RNA-seq

**DOI:** 10.1101/762773

**Authors:** Valentine Svensson, Eduardo da Veiga Beltrame, Lior Pachter

## Abstract

The allocation of a sequencing budget when designing single cell RNA-seq experiments requires consideration of the tradeoff between number of cells sequenced and the read depth per cell. One approach to the problem is to perform a power analysis for a univariate objective such as differential expression. However, many of the goals of single-cell analysis requires consideration of the multivariate structure of gene expression, such as clustering. We introduce an approach to quantifying the impact of sequencing depth and cell number on the estimation of a multivariate generative model for gene expression that is based on error analysis in the framework of a variational autoencoder. We find that at shallow depths, the marginal benefit of deeper sequencing per cell significantly outweighs the benefit of increased cell numbers. Above about 15,000 reads per cell the benefit of increased sequencing depth is minor. Code for the workflow reproducing the results of the paper is available at https://github.com/pachterlab/SBP_2019/.

## Introduction

The design of single-cell RNA-seq experiments requires numerous choices including allocation of a budget for sequencing, determination of how many cells will be assayed, the technologies to be used, and choices on how to extract and prepare cells. When the source of cells is plentiful, and there is a fixed budget for a single-cell RNA-seq (scRNA-seq) experiment, the key choice is whether to aim for an experiment with more cells and fewer reads per cell, or less cells but each cell sequenced more deeply. An analysis of this important tradeoff has been investigated in the context of learning parameters for single genes, and it was shown that more cells are preferable to higher read depth per cell [1].

Furthermore, for typical univariate supervised inference problems such as regression, tools for power analysis exist to assess what parameter values in models can be reliably inferred from data of varying sizes [2,3]. However, questions of interest in single-cell RNA-seq analysis do not have a structure where performance can be directly quantified and appropriate subsampling and quantification of performance is nontrivial. While there has been some work examining the dependence of principal components analysis on read depth [4,5], the methods used do not extend quantitatively to other multivariate analyses.

Key analysis tasks in scRNA-seq studies such as cell type discovery and assignment, or trajectory inference, make use of low-dimensional representations of the cells. As a result a number of generative models that identify low-dimensional representations have been proposed [6–8]. One recent development is the introduction of generative models with hidden representations in the form of variational autoencoders [9–13]. Since variational autoencoders are parametric they can easily be used to evaluate performance on held-out unseen validation data using the comparable and quantitative measure of log likelihood: the probability of seeing the data given the trained model (see methods). Negative log likelihood is also referred to as *reconstruction error* and here we refer to the reconstruction error of the held-out validation data as *validation error*.

## Results

To investigate the effect of sequencing depth and cell numbers on the ability to learn reproducible representations of scRNA-seq data, we performed a subsampling analysis of three datasets produced with the 10x Genomics Chromium platform (V3) [14]. The datasets were selected for their high sequencing depth and each dataset consisted of approximately 10,000 cells (**Supplementary Table 1**).

The kallisto bustools workflow [15–17] was used to obtain UMI [18] gene count matrices at different sampled read depths to mimic datasets sequenced at varying depths. From these subsamples, subsets of cells were sampled to emulate the sequencing of fewer cells **(Figure 1)**.

**Figure 1).**
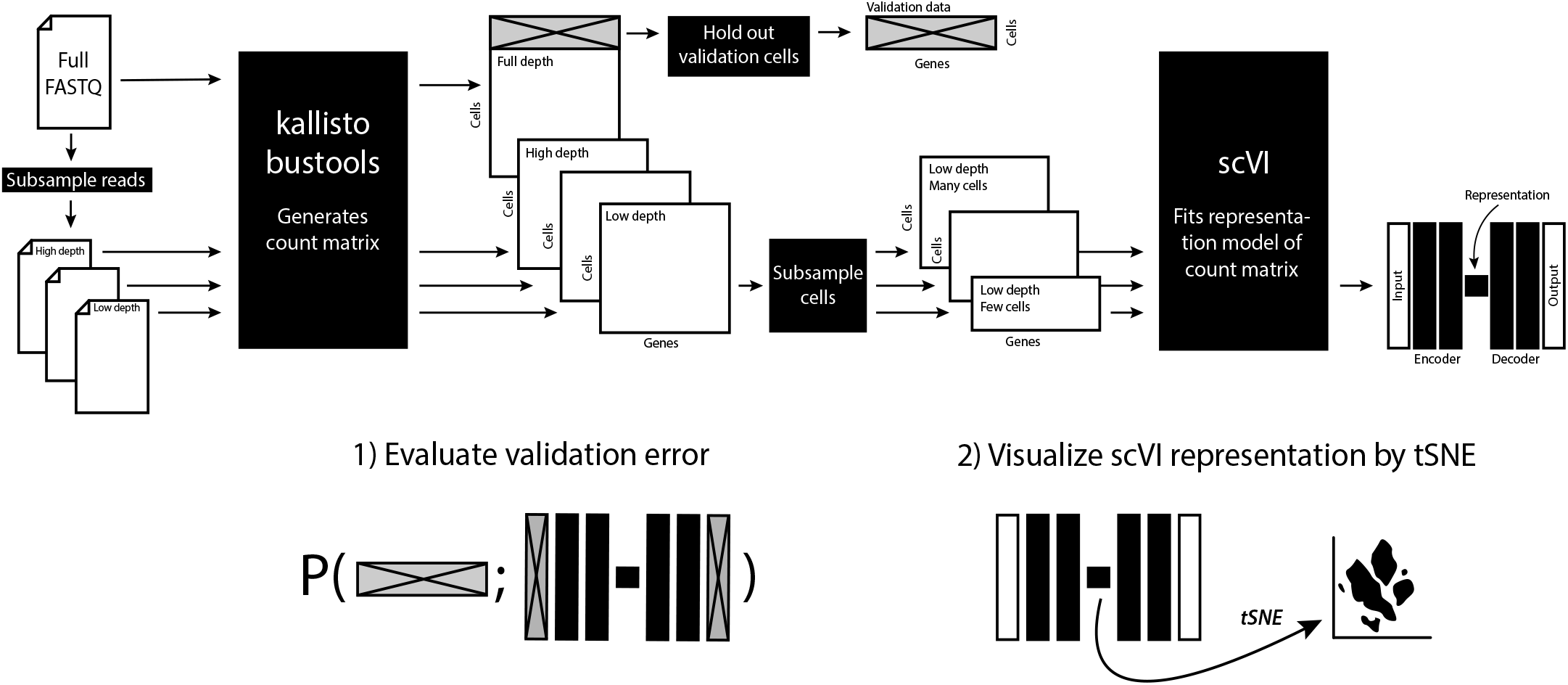
Outline of the workflow for subsampling reads and cells, fitting models with a variational autoencoder, evaluating validation error, and visualization.

For each of the gene count matrices with different reads per cell and cell numbers, an scVI variational autoencoder model was fit. A set of cells were held-out prior to subsampling and for later use as a validation set. Each of the scVI models were applied to the held-out data, and reconstruction error was calculated to give validation error values for each point in the sampling grid of cell numbers and reads per cell numbers **(Figure 2)**.

**Figure 2).**
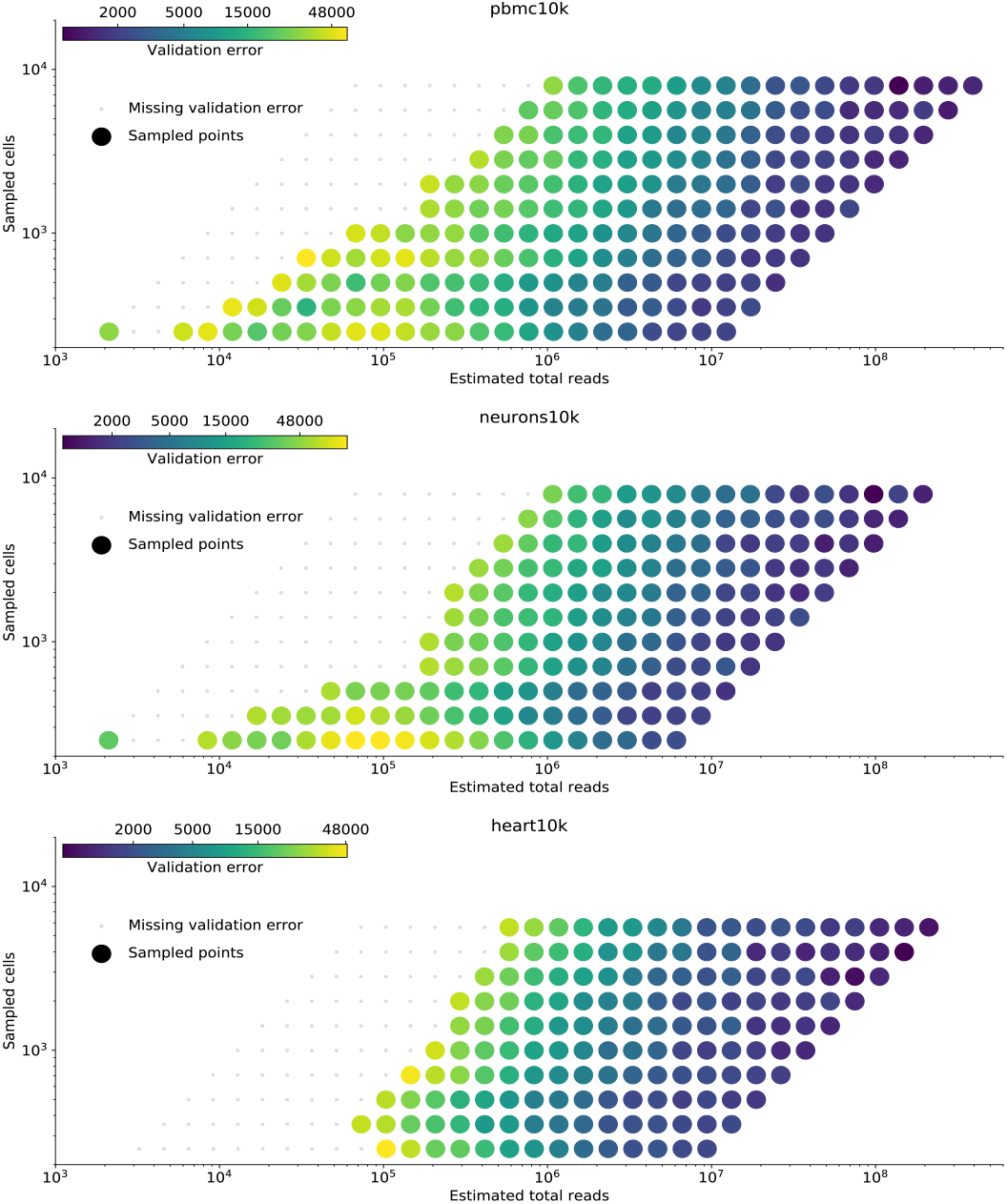
Held-out log likelihood for each read sampling depth and cell sampling number for all three datasets. Gray dots indicate validation error could not be evaluated due to division by zero errors.

We found that there exists an inflection point below which increasing sequencing depth rapidly reduces validation error. When there are less than 15,000 reads per cell, doubling the sequencing depth decreases the error by 35-40%. This can be directly compared to doubling the number of cells, which decreases validation error by only 10-15%. The advantage of more sequencing in the shallow sequencing regime is much greater than the advantage realized with assaying more cells. This trend tapers off with increasing sequencing depth: with more than 15,000 reads per cell, doubling the number of cells or reads offers comparable marginal improvement, with error decreasing by at most 1-3% (**Table 1**, **Figure 3**).

**Table 1).** Effect on held-out validation error when doubling number of cells or doubling number of reads per cell for the three datasets investigated. Effects stratified by when cells have below 15,000 reads per cell or above. A larger decrease in error is better.

|  | Below 15,000 reads per cell |  | Above 15,000 reads per cell |  |
| --- | --- | --- | --- | --- |
|  | Doubling number of cells | Doubling reads per cell | Doubling number of cells | Doubling reads per cell |
| **heart10k**<br>(decrease in validation error) | 15% | 39% | 1% | 2% |
| **neurons10k**<br>(decrease in validation error) | 16% | 41% | 3% | 2% |
| **pbmc10k**<br>(decrease in validation error) | 10% | 35% | 2% | 5% |

**Figure 3).**
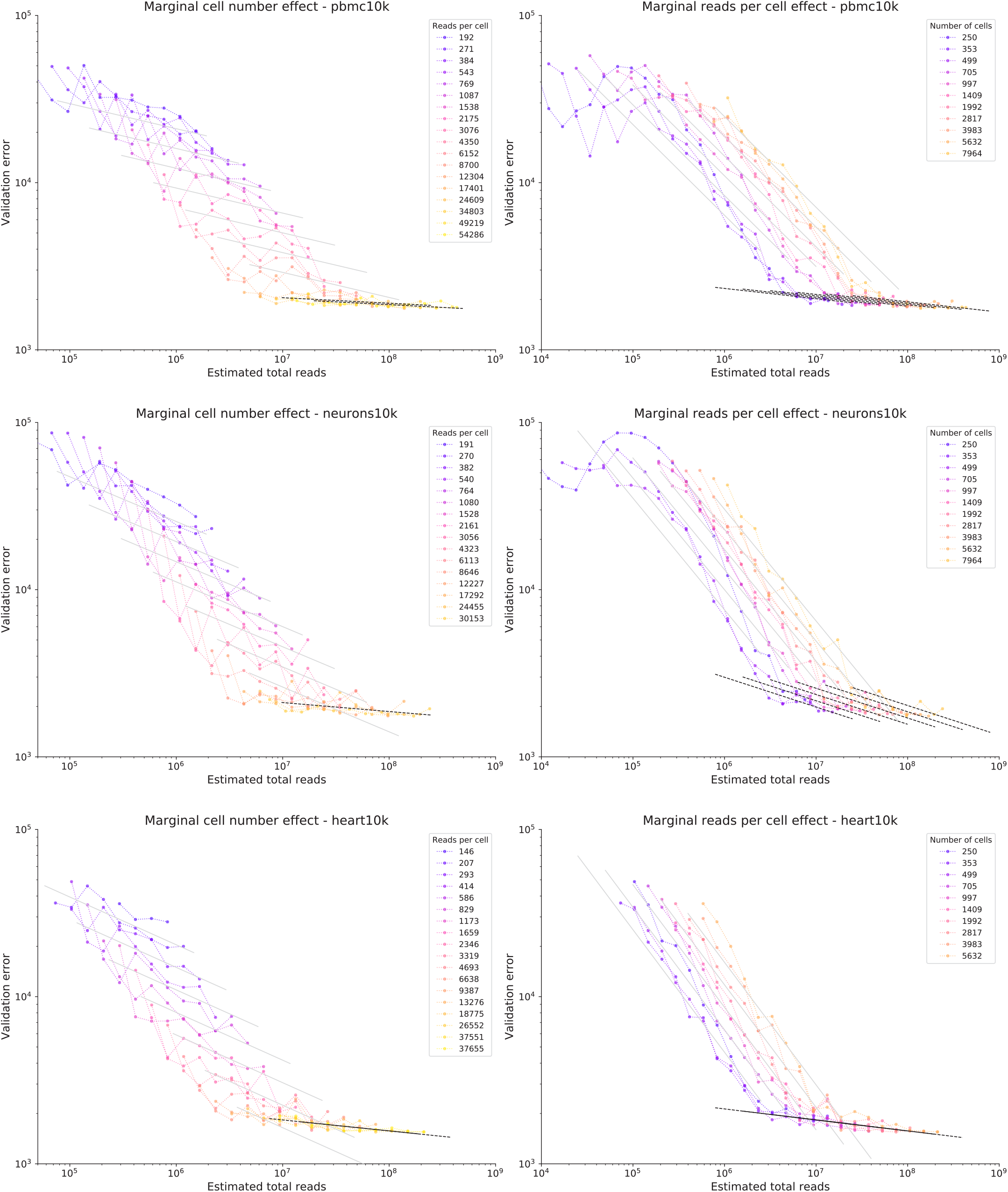
Validation error as a function of total reads. (**left**) Datasets grouped by reads per cell. (**right**) Datasets grouped by number of cells. (**top**) pbmc10k data, (**middle**) neurons10k data, (**bottom**) heart10k data. Grey and black lines are predictions from linear regression model, grey for < 15,000 reads per cell, black for >= 15,000 reads per cell. Each line is the prediction on a corresponding group of data.

Reconstruction error of the held out validation sets allow for quan titative analysis of the performance of models trained with different sets of data. To assess the implication of reconstruction error for downstream analyses, we performed t-SNE on the low-dimensional representation encoded by the scVI models for each read depth and cell sampling size. While the resultant plots are subjective in nature, the grid of t-SNE plots provides intuition for the loss of structure in the data corresponding to varying levels of reconstruction error. For all datasets, groupings of cells appear more distinct as read and cell numbers increase **(Figure 4)**.

**Figure 2).**
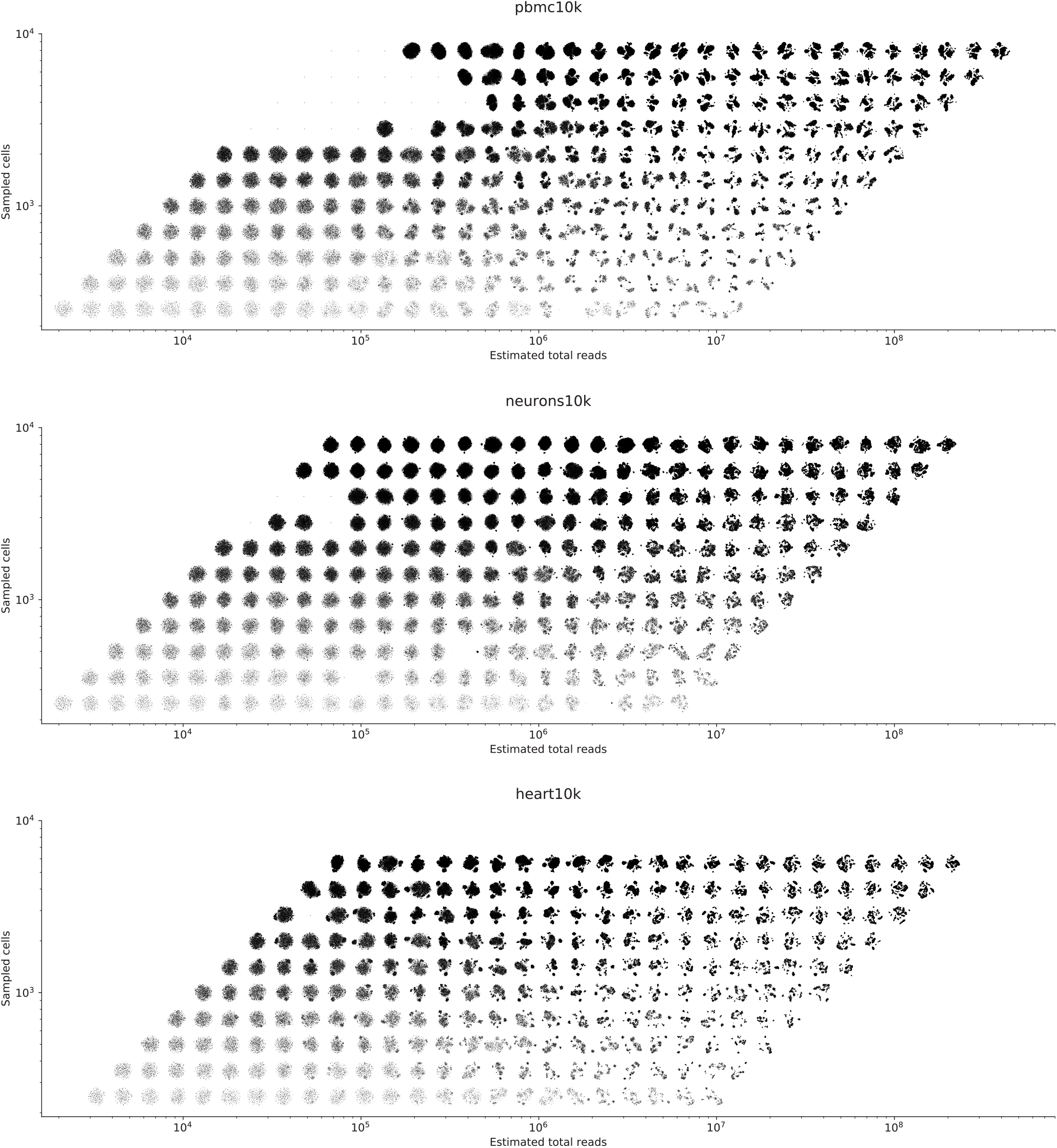
t-SNE visualization of the learned scVI representation for each read depth and cell count number for all three datasets. Plots with single dots correspond to representations where t-SNE failed to optimize due to small gradients. (**top**) pbmc10k data, (**middle**) neurons10k data, (**bottom**) heart10k data.

## Discussion

In this work we introduced the concept of autoencoder based “power analysis”. We implemented it to evaluate how sequencing depth and cell numbers affect the ability to learn generalizable representation models for single cell RNA-seq data. We found that increasing depth up to about 15,000 reads per cell has a substantially higher marginal benefit than increasing the number of cells. Beyond that, increasing either depth or cell numbers provides comparable marginal improvement. This constitutes a quantitative description of the tradeoff between sequencing depth per cell, and number of cells assayed.

We note that different protocols may have different efficiency in converting reads to UMIs due to different levels of PCR duplication. This may affect the slopes of marginal gain for number of cells or reads per cell. In the data used here, the duplication rate was consistently around 2.5 (**Supplementary Figure 1**). Examining the relationship between reads per cell and validation error with UMIs per cell and validation error, we found that results are identical up to multiplication by the duplication rate (**Supplementary Figures 2-4**). We note that this analysis does not take into account other experimental objectives that may be important. For example, if the goal of an experiment is to capture and characterize rare cells, the number of cells assayed should take into account the probability of capturing these cells of interest.

Our study yields some insights in terms of experimental design strategies utilizing a machine learning method. This approach may be also be productive for other tasks where tradeoffs in data generation cost must be considered. Our concept of autoencoder based power analysis using held-out log likelihood on augmented data may therefore be of use for other problems in learning representations. Some examples are machine learning models for text-to-speech, image recognition problems, or biological sequence data. Our workflow is available at https://github.com/pachterlab/SBP_2019/ and can be readily applied to any other scRNA-seq dataset that can be processed using kallisto bus (https://kallistobus.tools).

## Methods

### Data

Three datasets made available by 10x Genomics were analyzed: pbmc10k, heart10k and neurons10k. They are summarized in (**Supplementary Table 1**).

### Training data evaluation using held-out log likelihood

In typical study design, power calculations are used to estimate the amount of required sampling for satisfactory type II error in a hypothesis testing framework. In many situations, it may also be important to assess the extent and nature of sampling needed to accurately estimate parameters for a (pre-registered) statistical model, or the data needed to select a model. Such calculations have direct bearing on the budget needed to perform the study.

One approach to quantifying accuracy when fitting or selecting models is to estimate goodness-of-fit by performing holdout validation (also known as out-of-sample validation). With this approach, a dataset is divided into two: a training set, and separately a validation set, which is held-out during training procedures and used only to assess results after parameters have been estimated.

Formally, we denote the dataset by *X*, and partition it into a training set 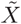 and a validation set 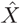. The latter is held-out from all processes of estimating a generative model 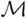 with parameters *θ*. The log-likelihood of held-out data is defined as the integral

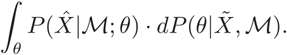

Here 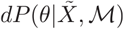 is the posterior probability of parameters of the model 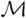 after it has been fitted with 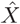. When point estimates of *θ* are considered, we have that

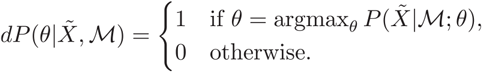

If *θ*\* is the maximum likelihood point estimate of *θ* for 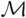 given 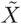 then the integral above reduces to 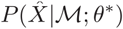.

By calculating the held-out log likelihood for two alternative models 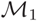 and 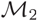, the models can be directly compared. In the publication describing scVI ([9]), this approach is used to compare the models of scVI with those obtained by factor analysis (principal component analysis), ZIFA [7], and ZINB-WaVE [19].

We use the same strategy to evaluate different sources of *training data* 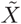. A fixed set of validation data 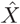 is pre-determined, and *K* different alternate data samples 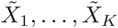 are generated. For example, the data could be generated from different annotation services, different scientific instruments, or with different sampling strategies. The model 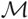 is fixed, but for each alternate training data set 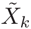, a maximum likelihood parameter estimate 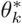 is estimated. This allows for the comparison of *K* different goodness-of-fits 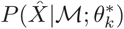.

A simple instance of this procedure that is of interest in virtually any modeling setting is the question of the generalizability of a model as a function of the amount of data observed. By subsampling data points from 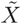 to different levels 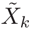, the return on investment from increased sample sizes can be examined using the directly comparable 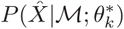 likelihoods. This provides a tool for study design that can be applied in the same way as power and sample size calculations are used in classical statistical analysis. Generating data for models can be expensive, and it is not uncommon to have a choice between different sources of data at different prices. Thus, this framework allows researchers to investigate how a data collection budget can be efficiently allocated.

### Subsampling scRNA-seq data

The result of a scRNA-seq experiment is a cDNA library which contains multiple copies of labeled mRNA transcript fragments, allowing for counting and identification of cell of origin. The count of molecules for each gene and cell make up a gene count matrix *X*.

For sequencing, the cDNA library is sampled and the molecules in that sample are distributed over an imaging array which is used to read out the DNA sequence of amplified products of each molecule. The density and size of the array determines how many “reads” the DNA sequencing instrument will deliver. It is possible (and common) to take multiple samples from the cDNA library which get sequenced on multiple imaging arrays or even different instruments. This allows fine control over the total depth (total number of reads) of the resulting data that is used to obtain the gene count matrix.

To simulate sequencing a cDNA library at a lower depth, randomly subsampling reads from the output of a DNA sequencer faithfully simulates generating data with a lower density imaging array, or using fewer imaging arrays. For a given simulated depth, after the gene count matrix with all cells was obtained, a dataset emulating the input of fewer cells can be generated by randomly sampling cells, which are treated as observations.

By first varying the ‘*reads per cell*’ through subsampling the list of reads, then subsampling cells conditional on this, a collection of alternative training datasets 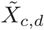 representing different resource allocation strategies can be created. Here refers to the number of cells and the ‘*reads per cell*’ in each dataset. The sampled cells are excluding a fixed set of validation cells making up the validation gene count matrix 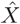.

### scVI variational autoencoder

The variational autoencoder implementation in scVI parametrizes a negative binomial distribution for each gene *g* in a cell *i* based on a low dimensional latent representation *z*_*i*_ which is “decoded” as *m*_*i*_ = *f*_*θ*_(*z*_*i*_). The generative model of a gene count matrix *X* is

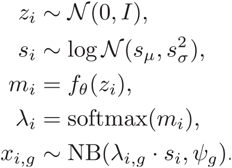

In this framework *s*_*i*_ is a scaling factor for the read depth and *m*_*i*_ is a vector of *scale* values for the gene expression levels which are transformed to the count rates λ_*i*_. In scVI, amortized inference is used to learn variational distributions *Q*(*z*|*x*) approximating the posterior distribution *P*(*z*_*i*_|*X*) using a pair of neural networks *g* parametrized by *ϕ* = {*ϕ*_*μ*_, *ϕ*_*σ*_}:

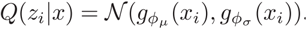

We refer to the full scVI model including its variational distributions as 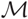 with parameter set Θ = {*θ*, *ϕ*}.

Parameters *θ* in an scVI variational autoencoder model 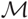, and variational encoder parameters *ϕ*, are found through optimizing the (approximate) marginal likelihood using a sample of data 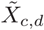 of desired *reads per cell d* and number of cells from training sets:

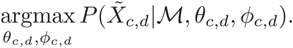

A held-out validation dataset 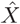 is then used to calculate the held-out marginal log likelihood, which allows direct comparison between alternative models, in this case differing by fitted values for *θ* and *ϕ*,

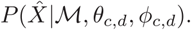

The approximate marginal likelihood of the scVI autoencoder is calculated by first sampling *Z* from the variational distribution of the representation, then decoding from each *z*_*i*_ the parameters λ_*i*,*g*_.

Finally the approximate marginal log likelihood is computed using

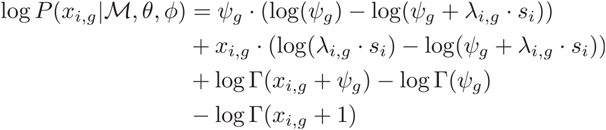

where

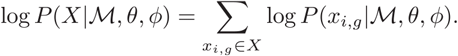

### Workflow description

The fastq files were subsampled at equal (multiplicatively) spaced intervals starting with 100,000 reads, and going up by a factor of √2 for each sampled point until the total number of reads in the dataset was reached. FASTQ subsampling was done using seqtk v1.3 (https://github.com/lh3/seqtk) with the random seed = 100. For each subsample, UMI count matrices were produced using kallisto v0.46.0 (https://pachterlab.github.io/kallisto/) and bus-tools v0.39.3 (https://bustools.github.io). Ensembl release 96 was used for the mouse and human transcriptomes and annotation (https://uswest.ensembl.org/info/data/ftp/index.html).

The scVI program (https://github.com/YosefLab/scVI) was used to create a 10-dimensional representation for each dataset using two hidden layers, each with 128 nodes. Negative binomial reconstruction loss was used to ascertain error. A set of 1,000 highly variable genes were determined using the full dataset, and these genes were used for each subsampled dataset. For each model, training was done with an 80% train-test ratio. The number of epochs was set to 27 * 10000 / (number of cells). This training rule was used based on empirical evidence of the convergence behavior of scVI.

Before subsampling cells, a random set of cells that were quantified without subsampling reads were removed from the dataset (validation set): 1,770 out of 11,769 for PBMCs (15%); 1,844 out of 11,843 for neurons (16%); 714 out of 7,713 for heart (9%). These cells were excluded from all subsampled count matrices before sampling cells and training the models.

Linear regression was performed on log2(validation error) using log2(number of cells) and log2(reads per cell) as covariates with the statsmodels package v0.9.0. Separate models were fit for points having reads per cell above and below 15,000. Evaluating different thresholds from 15,000 resulted in similar effect sizes for number of cells and reads per cell (**Supplementary Table 2**).

Each of the 10-dimensional representations learned by scVI for the different subsampled data was visualized using t-SNE. This was performed using the openTSNE package v0.3.10 (https://github.com/pavlin-policar/openTSNE) with random initialization, approximate nearest neighbor calculation, FFT based negative gradient calculations, and was run for 1,000 iterations with the default perplexity parameter of 30.

## Code availability

A Snakemake [20] file used to subsample and process the data, together with Python notebooks used for downstream analyses are available on GitHub at https://github.com/pachterlab/SBP_2019/. Scripts and notebooks used to create the figures and results, together with gene count matrices outputted by kallisto bus and H5AD files with the UMI counts for all the subsampled read depths are available on CaltechDATA (https://doi.org/10.22002/d1.1276).

## Author contributions

V.S. designed the evaluation metric and performed statistical analysis. E.V.B. performed data processing and subsampling. V.S., E.V.B., and L.P. interpreted results and wrote the manuscript.

## Acknowledgements

The authors want to thank Romain Lopez for helpful feedback on the manuscript. V.S. and L.P. were funded in part by NIH U19MH114830.

## Supplemental material

**Supplementary Figure 1).**
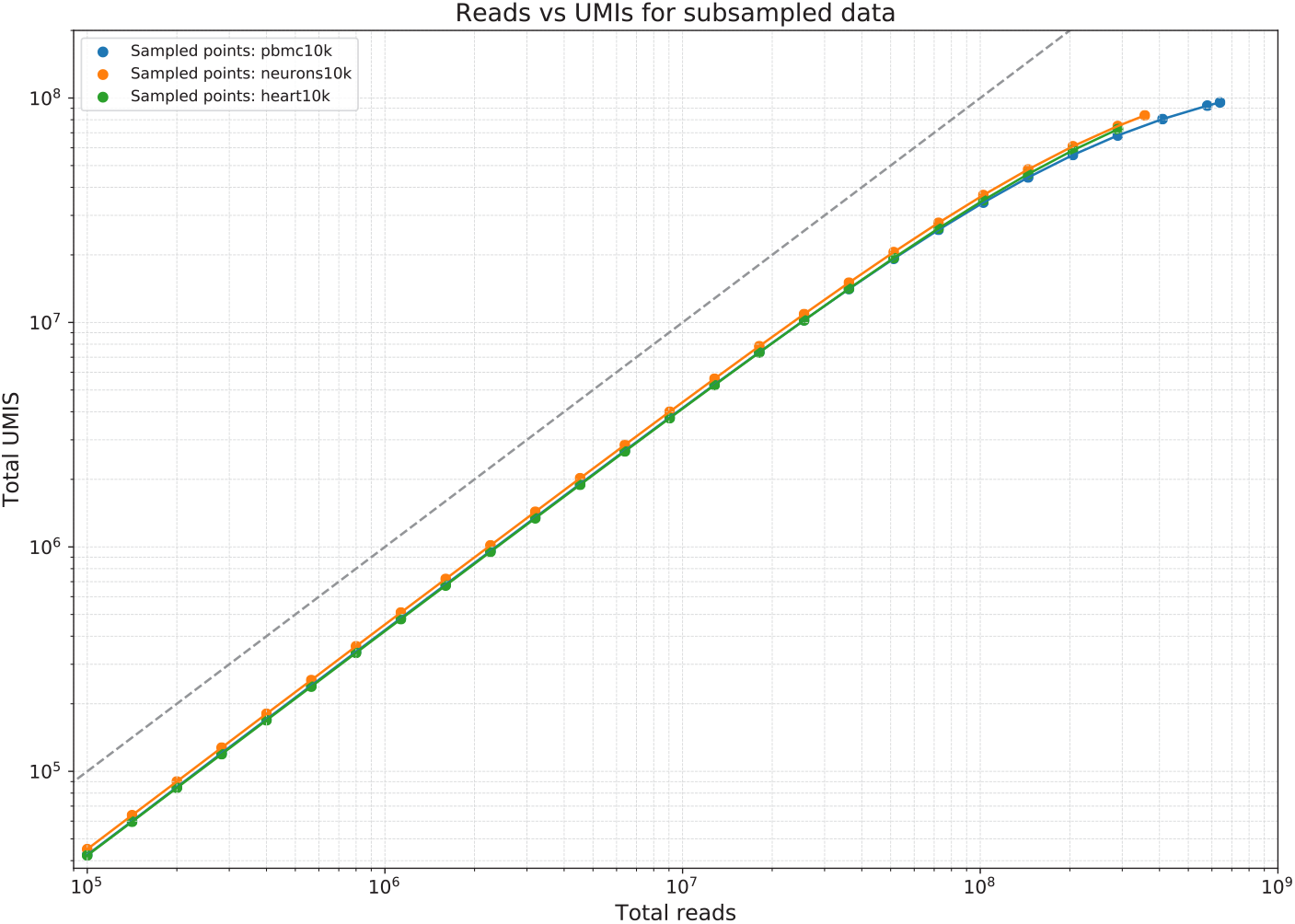
Reads vs UMIs for all datasets at subsampled depths.

**Supplementary Figure 2).**
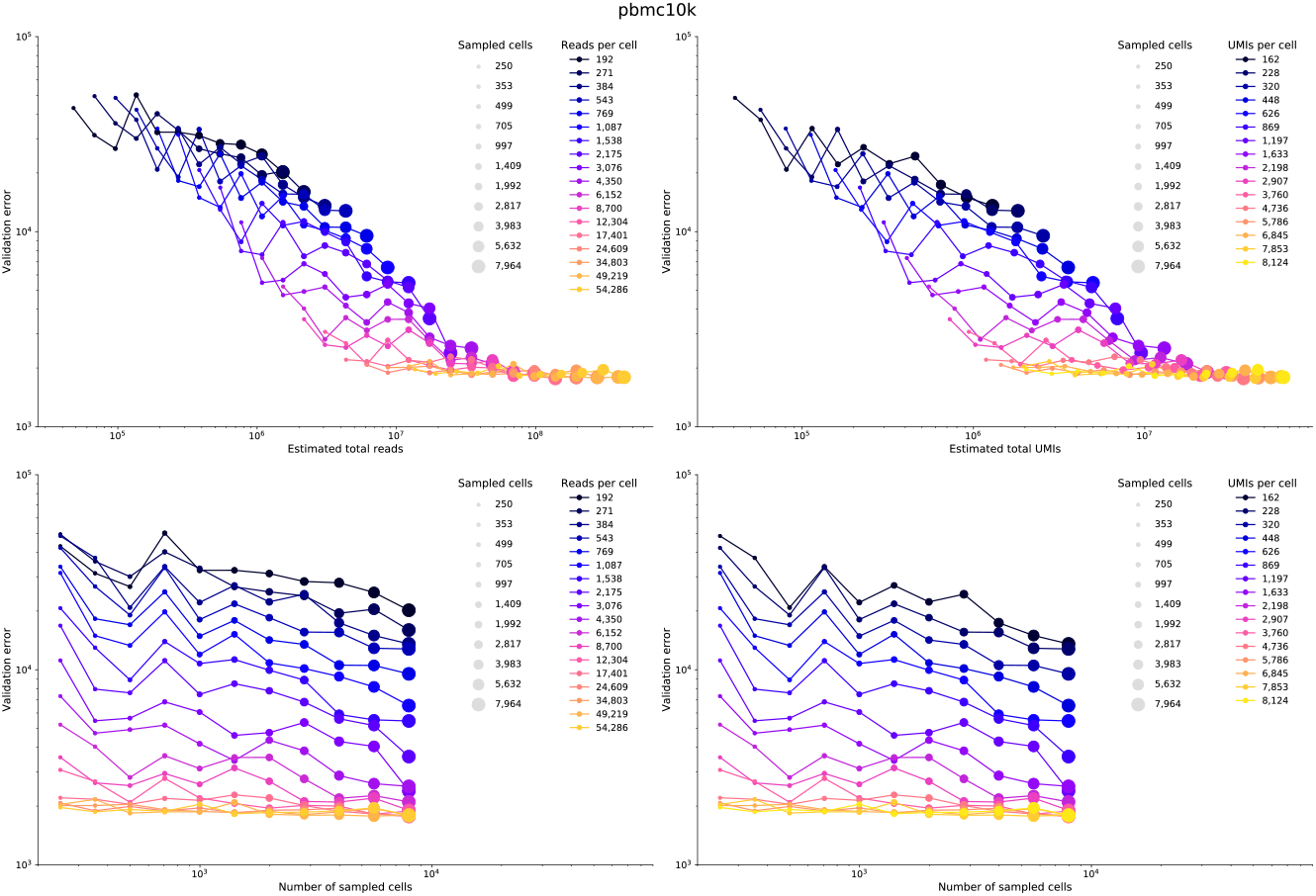
pbmc10k composition. **Top left**: Total reads vs validation error. **Top right**: Total UMIs vs validation error. **Bottom left**: Sampled cells vs validation error, plotted over reads. **Bottom right**: Sampled cells vs validation error, plotted over UMIs.

**Supplementary Figure 3).**
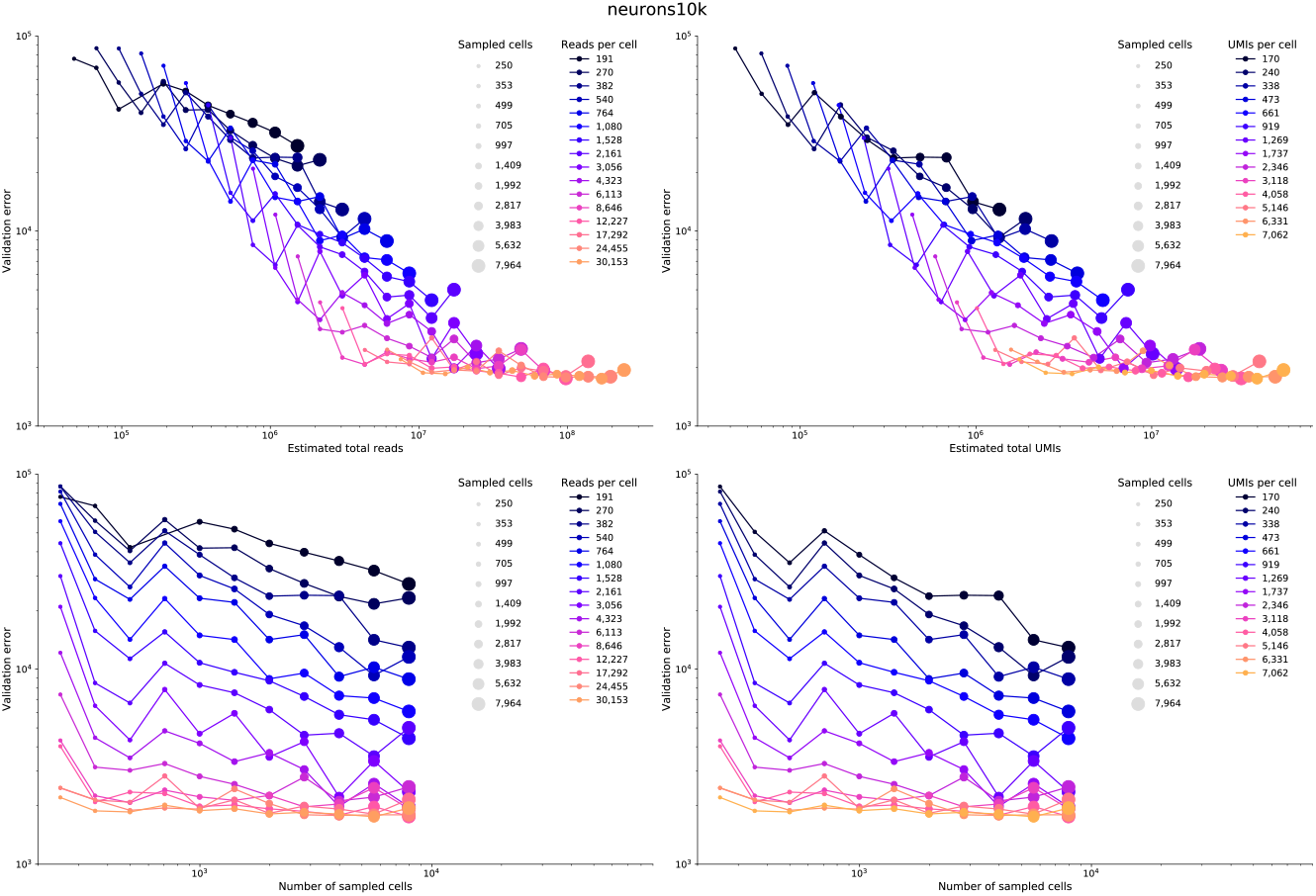
neurons10k composition. **Top left**: Total reads vs validation error. **Top right**: Total UMIs vs validation error. **Bottom left**: Sampled cells vs validation error, plotted over reads. **Bottom right**: Sampled cells vs validation error, plotted over UMIs.

**Supplementary Figure 4).**
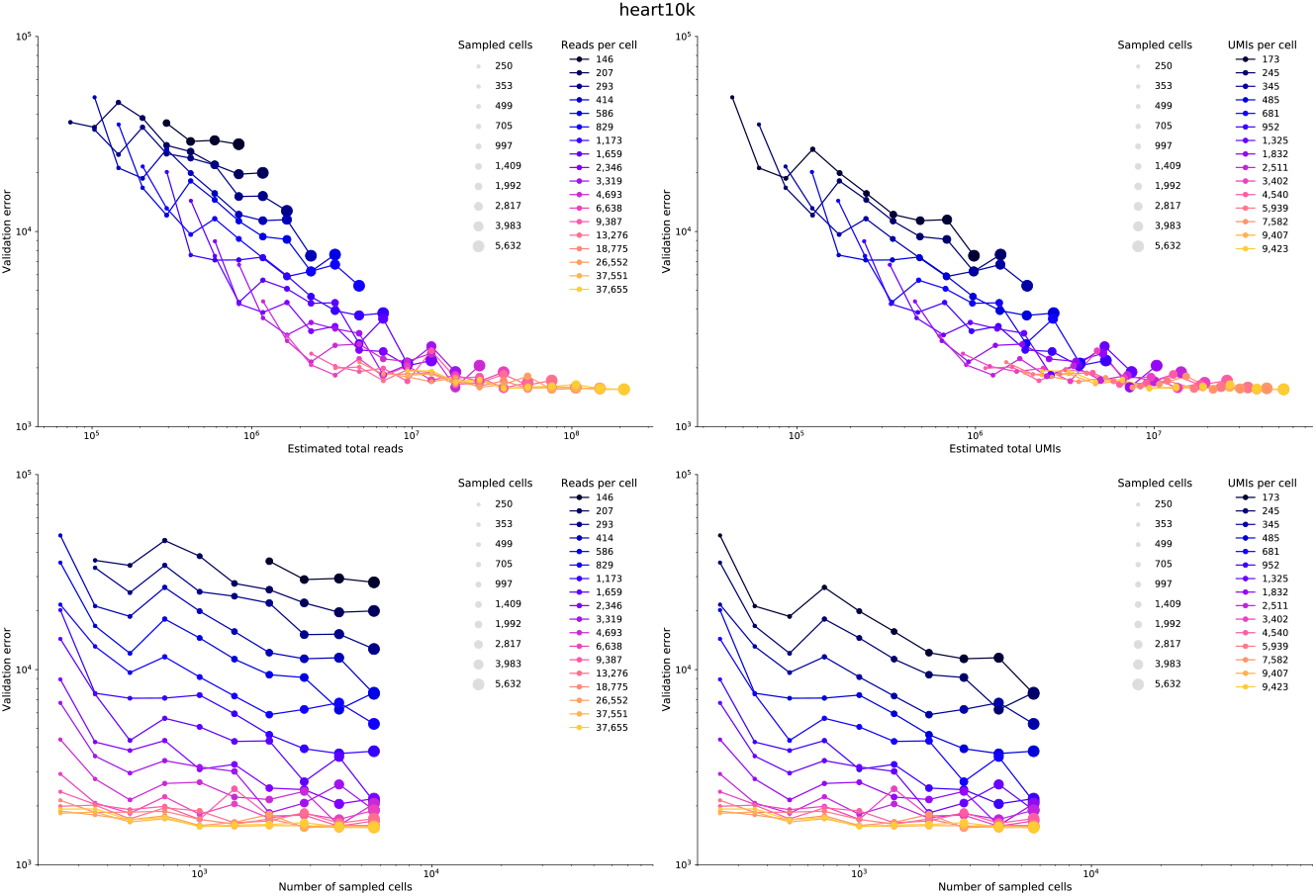
heart10k composition. **Top left**: Total reads vs validation error. **Top right**: Total UMIs vs validation error. **Bottom left**: Sampled cells vs validation error, plotted over reads. **Bottom right**: Sampled cells vs validation error, plotted over UMIs.

**Supplementary Table 1).**
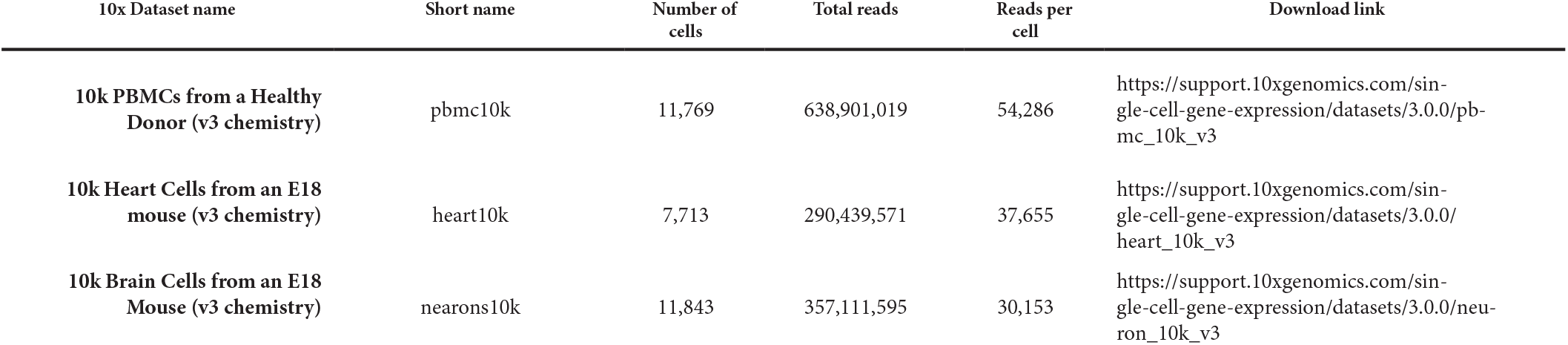
Summary of the three datasets analyzed.

**Supplementary Table 2).**
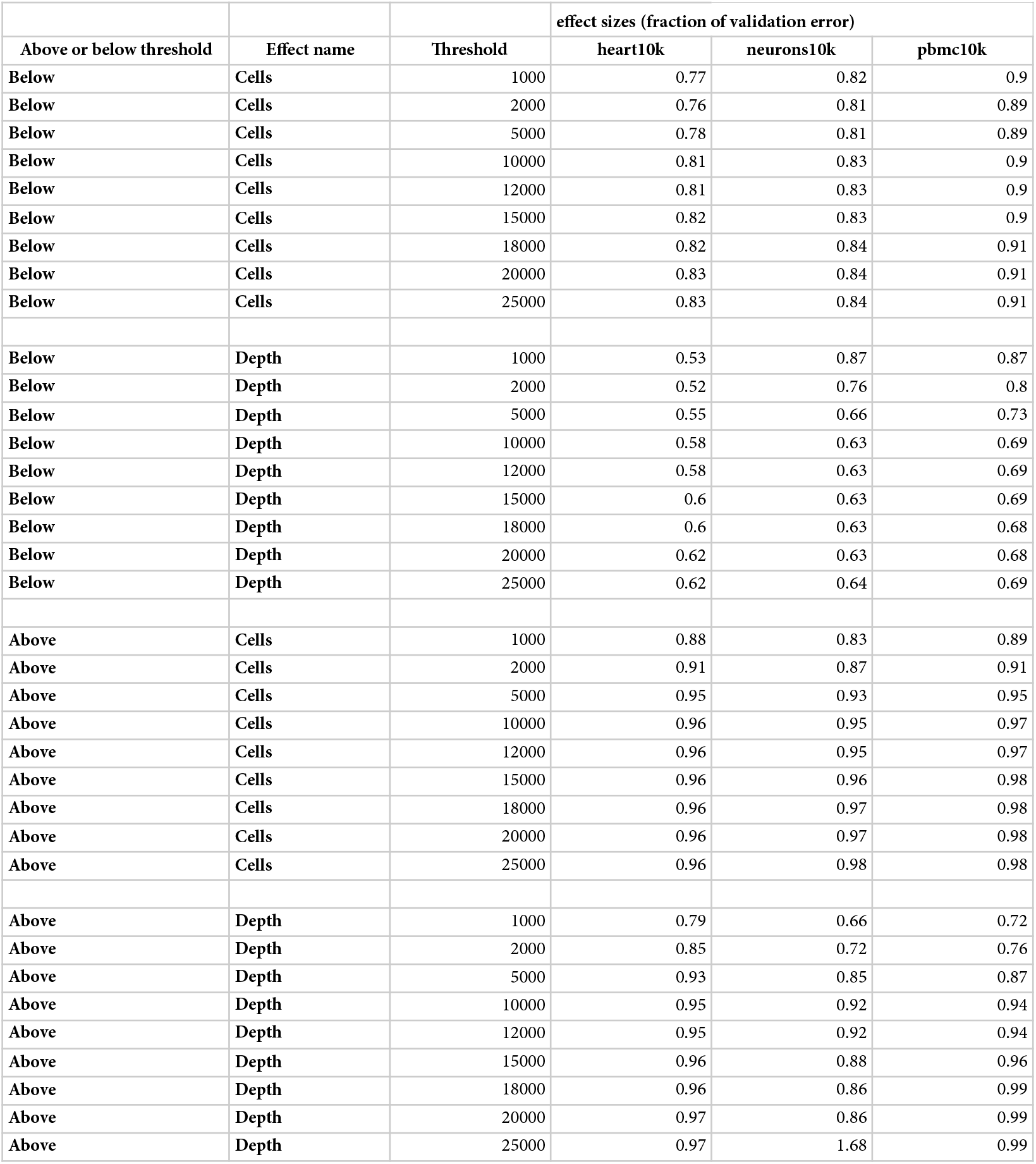
Effect sizes for number of cells and reads per cells for different values of the reads per cell threshold for the three datasets.

## References

1. Zhang MJ, Ntranos V, Tse D. One read per cell per gene is optimal for single-cell RNA-Seq. 2018; 15.

2. Vieth B, Ziegenhain C, Parekh S, Enard W, Hellmann I. powsimR: power analysis for bulk and single cell RNA-seq experiments. Bioinformatics. 2017;33: 3486–3488.

3. Bass AJ, Robinson DG, Storey JD. Determining sufficient sequencing depth in RNA-Seq differential expression studies [Internet]. bioRxiv. 2019. p. 635623. doi:10.1101/635623

4. Heimberg G, Bhatnagar R, El-Samad H, Thomson M. Low Dimensionality in Gene Expression Data Enables the Accurate Extraction of Transcriptional Programs from Shallow Sequencing. Cell Syst. 2016;2: 239–250.

5. Ntranos V, Kamath GM, Zhang JM, Pachter L, Tse DN. Fast and accurate single-cell RNA-seq analysis by clustering of transcript-compatibility counts. Genome Biol. 2016;17: 112.

6. Risso D, Perraudeau F, Gribkova S, Dudoit S, Vert J-P. A general and flexible method for signal extraction from single-cell RNA-seq data. Nat Commun. 2018;9: 284.

7. Pierson E, Yau C. ZIFA: Dimensionality reduction for zero-inflated single-cell gene expression analysis. Genome Biol. 2015;16: 241.

8. Azizi E, Carr AJ, Plitas G, Cornish AE, Konopacki C, Prabhakaran S, et al. Single-Cell Map of Diverse Immune Phenotypes in the Breast Tumor Microenvironment. Cell. 2018;174: 1293–1308.e36.

9. Lopez R, Regier J, Cole MB, Jordan MI, Yosef N. Deep generative modeling for single-cell transcriptomics. Nat Methods. 2018;15: 1053–1058.

10. Lotfollahi M, Wolf FA, Theis FJ. scGen predicts single-cell perturbation responses. Nat Methods. 2019;16: 715–721.

11. Grønbech CH, Vording MF, Timshel PN, Sønderby CK, Pers TH, Winther O. sc-VAE: Variational auto-encoders for single-cell gene expression data [Internet]. bioRxiv. 2018. p. 318295. doi:10.1101/318295

12. Ding J, Condon A, Shah SP. Interpretable dimensionality reduction of single cell transcriptome data with deep generative models. Nat Commun. 2018;9: 2002.

13. Ghahramani A, Watt FM, Luscombe NM. Generative adversarial networks simulate gene expression and predict perturbations in single cells [Internet]. bioRxiv. 2018. p. 262501. doi:10.1101/262501

14. Zheng GXY, Terry JM, Belgrader P, Ryvkin P, Bent ZW, Wilson R, et al. Massively parallel digital transcriptional profiling of single cells. Nat Commun. 2017;8: 14049.

15. Melsted P, Sina Booeshaghi A, Gao F, Beltrame E, Lu L, Hjorleifsson KE, et al. Modular and efficient pre-processing of single-cell RNA-seq [Internet]. bioRxiv. 2019. p. 673285. doi:10.1101/673285

16. Bray NL, Pimentel H, Melsted P, Pachter L. Near-optimal probabilistic RNA-seq quantification. Nat Biotechnol. 2016;34: 525–527.

17. Melsted P, Ntranos V, Pachter L. The Barcode, UMI, Set format and BUStools. Bioinformatics. 2019; doi:10.1093/bioinformatics/btz279

18. Kivioja T, Vähärautio A, Karlsson K, Bonke M, Enge M, Linnarsson S, et al. Counting absolute numbers of molecules using unique molecular identifiers. Nat Methods. 2011;9: 72–74.

19. Risso D, Perraudeau F, Gribkova S, Dudoit S, Vert J-P. Publisher Correction: A general and flexible method for signal extraction from single-cell RNA-seq data. Nat Commun. 2019;10: 646.

20. Köster J, Rahmann S. Snakemake-a scalable bioinformatics workflow engine. Bioinformatics. 2018;34: 3600.

